# Visualization of neuronal morphology of the suprachiasmatic nucleus vasopressin neurons by Cre/FLPe-based genetic two-step sparse labelling—GT-SPARCL

**DOI:** 10.64898/2026.05.25.727737

**Authors:** Sui-Wen Hsiao, Yoshiaki Yamaguchi, Daichi Moriyasu, Shotaro Doi, Huihua Zhou, Seiya Mizuno, Satoru Takahashi, Hiroko Yukinaga, Tom Macpherson, Emi Hasegawa, Masao Doi

**Affiliations:** Department of Systems Biology, Graduate School of Pharmaceutical Sciences, Kyoto University, Sakyō-ku, Kyoto 606-8501, Japan; Laboratory Animal Resource Center in Transborder Medical Research Center, Institute of Medicine, University of Tsukuba, Tsukuba, Ibaraki 305-8575, Japan; Department of Life Science, Graduate School of Life Science, University of Hyogo, Ako-gun, Hyogo 678-1297, Japan

**Keywords:** Sparse labelling, Suprachiasmatic nucleus, Arginine vasopressin neurons

## Abstract

Resolving the morphology of individual neurons in densely packed brain regions remains challenging. Sparse labeling is essential for delineating cellular architecture, yet achieving reproducible low-density labeling e.g. < 1% has been a persistent technical hurdle/problem. We present a genetically encoded two-step strategy, GT-SPARCL (Genetic Two-Step Sparse Cre/FLPe Combination Labeling), which leverages two independent stochastic recombination events to deliver stable, tunable low-density labeling in mice. As a proof of concept, we applied this method to visualizing arginine vasopressin (AVP) neurons in the suprachiasmatic nucleus (SCN), the master circadian pacemaker composed of ∼10,000 neurons per side in mice. Using two-photon imaging of whole-mount SCN blocks, we reconstructed individual AVP neurons and uncovered previously under-resolved morphological heterogeneity. Based on axonal trajectories, we distinguished five structural types, including subclasses with commissural projections to the contralateral SCN and others extending projections beyond the nucleus. The majority (∼70%) exhibited projections both within and outside the ipsilateral SCN, whereas the second most dominant subset (∼20%) projected exclusively outside the SCN, representing an output-biased type. In contrast, neurons projecting exclusively within the ipsilateral SCN were exceptionally rare, suggesting that “dedicated” local-circuit AVP neurons do not form a major structural subtype. Collectively, our data indicate that AVP neurons are not structurally uniform but instead comprise diverse projection-defined subtypes, implying subtype-dependent contributions to intra-SCN communication, bilateral coupling, and circadian output. Beyond the SCN, our GT-SPARCL method may be applicable for achieving low-density labelling of neurons that can be defined by other specific Cre mouse lines.

## INTRODUCTION

The foundation of neuroscience begins with Cajal’s systematic analyses of Golgi-stained neuronal morphology in the vertebrate brain, which revealed the structural principles of individual neurons and their circuit formation^1,2^. However, several barriers still exist in continuing this endeavor. Neurons in the brain are densely packed and their intermingled axonal and dendritic processes obscure the morphology of individual cells. Sparse labeling is therefore indispensable for visualizing structures of specific neurons of interest^3,4^.

In sparse labelling, a few selective neurons should be labeled in a given population. However, neuronal populations are diverse in cell numbers and cell density. Therefore, appropriate sparse labelling must achieve an ideal low-density cell labelling ratio, ranging from as small as ∼0.01 % to 10% depending on the density and cell number of the neuronal cluster studied. Attaining a consistently low-rate labelling, e.g. below 0.1%, is however difficult (challenging) as it becomes statistically variable in labelling reaction. The traditional Golgi method exhibit a labelling ratio of 1-5%. Virus-based methods also display a range of diluted (0.1%−10%) labelling ratios but are sensitive to variable infection efficiency. A recently developed genetic mouse-based sparse labelling method, MORF (ref^2^), has a labelling ratio from 1% to 5% depending on the type of Cre mouse line used. Thus, there is still room for improvement in achieving sparse labelling at ratios smaller than 1% or 0.1%. To mitigate this problem, we sought to develop a genetic based “two-step” sparse labelling method. The core concept of this method lies in combining two independent stochastic events in sequence, which together yield a deeper level of sparseness. More specifically, we established two competitive (stochastic) recombination systems by modifying Cre recombinase-loxP (lox3172) and Flippase (FLPe) recombinase-FRT (F3) systems.

Moreover, by introducing these two systems into mouse Rosa26 allele (genetic knock-in line), we aimed to obtain genetically stable lower labelling ratios—ideally below 1%— which may be helpful for analyzing neuronal clusters containing thousands of neurons.

The hypothalamic suprachiasmatic nucleus (SCN) is a pair of compact nuclei, each containing ∼10,000 tightly packed neurons. Physiologically, the SCN is the locus of the master circadian pacemaker, governing daily rhythms in behavior and diverse physiologies including core body temperature, sleep–wake cycle, metabolism, and hormonal secretion^5–7^. However, regarding the morphological information on SCN neurons, available data are limited and largely derived from traditional Golgi staining or manual biocytin labeling^8–10^. Although several recent connectomic studies involving sparse-labeling techniques have been published^11,12^, comparable datasets for the SCN are missing^13^, perhaps owing to its dense and compact anatomical architecture.

In the SCN, neurons expressing the neuropeptide arginine vasopressin (AVP) account for approximately 20% of the total neuronal population, corresponding to roughly 4,000 cells. Although AVP cell type-specific knockout studies (e.g., circadian clock gene knockout using AVP-Cre mouse line) elucidated the physiological importance of this cell type, morphological studies are currently limited^8,14^. Earlier studies based on Golgi staining provided pictures of SCN neurons, but this approach was not cell type-specific and cannot address AVP neurons^8^. On the other hand, studies utilizing biocytin labelling followed by anti-AVP immunostaining provided several examples of the morphology for AVP neurons^14^. These observations suggest that AVP neurons are morphologically heterogeneous, comprising neurons that only form local circuits inside the SCN as well as those possessing axonal projections to both inside and outside of the SCN^14^. Nevertheless, as the authors of these studies noted, axons extending beyond the plane of the SCN tissue sections were severed during sectioning, which might obscure the full morphology of individual AVP neurons in the SCN (in the whole mount SCN).

In our study, with the aim of developing a proof of concept of our two-step sparse labelling method, and to capture a previously uncharacterized morphology of SCN AVP neurons, we applied our genetic-based SPARCL system to visualize AVP neurons in the SCN.

## RESULTS

### Design and construction of SPARCL knock-in mice

The SPARCL strategy is schematically illustrated in Fig. 1. In order to achieve a stable sparse labelling of SCN neurons, we employed genetically-encoded two-step competitive (stochastic) recombination strategy, composed of SPARCL-FLPe (Fig. 1A) and SPARCL-ChR2-tdTomato/ Syp-EGFP (Fig. 1B and B’) Rosa26 genetic knock-in (KI) lines (see also STAR METHODS). For the former FLPe^KI^ line, two *lox* pairs—loxP and lox3172—were employed for competition: lox3172 is a loxP variant with reduced recombination efficiency^15^, adopted to reduce the probability of FLPe expression. On the other hand, loxP-dependent recombination was designed to result in E2Crimson expression to localize Cre^+^ cells. The loxP and lox3172 were arranged in a cross-linked configuration; this ensures mutually exclusive expression of FLPe and E2Crimson. The reduced recombination for FLPe-expression was further assisted by the distance between lox3172 sites (2.6 kb) that is larger than the length between loxP elements (0.9 kb) (Fig. 1A). In the latter SPARCL-ChR2-tdTomato/Syp-EGFP^KI^ line, we also devised a similar competing recombination strategy using two FLPe-recombination pairs, FRT and F3. F3 is a FRT variant with reduced recombination efficiency^16^, aiding to reduce the number of reporter gene (ChR2-tdTomato/Syp-EGFP)-expressing cells in the SCN. To further modulate the ratio (sparseness) of positive cell labelling, we elongated the length between F3 sites, based on the hypothesis that a greater distance between recombination sites would reduce recombination efficiency; thus, we constructed two mouse lines whose inter-F3 distance is either 2.6 kb or 16.3 kb due to the absence or presence of a 13.7-kb spacer sequence (Fig. 1B and B’). In both constructs, occurrence of F3 recombination results in initiation of reporter gene expression. Our reporter encodes P2A sequence-interlinked ChR2-fused tdTomato (ChR2-tdTomato) and synaptophysin -fused EGFP (Syp-EGFP): ChR2-tdTomato localizes to the cell membrane and helps visualize the entire soma and neurites of labeled neurons, whereas Syp-EGFP localizes to the presynaptic terminals and serves to visualize synapse locations^17,18^. Occurrence of recombination between FRT-sites, on the other hand, causes expression of E2Crimson (so that we can label all Cre^+^ cells examined). By integrating two sequential recombination events, namely Cre-dependent SPARCL and FLPe-dependent SPARCL, in a single mouse (i.e. dual positive heterozygous mouse), we aimed to perform sparse labelling based on genetically controlled mice.

**Figure 1.**
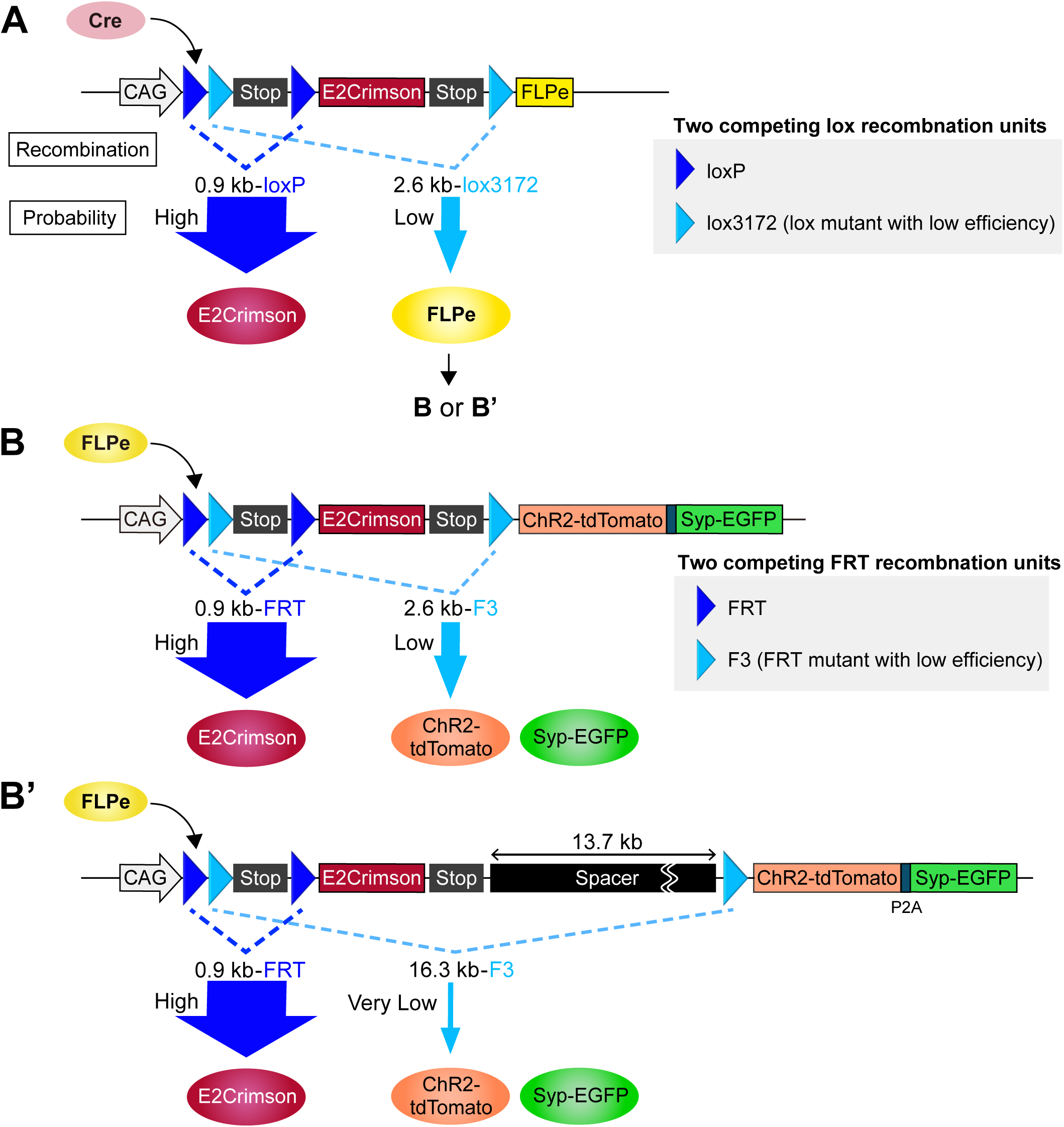
The two-step competitive recombination strategy in SPARCL. (A) Rosa26-SPARCL-FLPe knock-in locus for Cre recombination. Two different recombination units, loxP and lox3172, enable mutually exclusive expression of E2Crimson (high probability, dark blue) and FLPe (low probability, light blue). The latter expression (FLPe) allows the next step recombination, illustrated in (B)(B’). (B) (B’) Rosa26-SPARCL-ChR2-tdTomato/Syp-EGFP knock-in locus with or without spacer sequence for FLPe recombination. Two distinct FLPe-recombination units, FRT and F3, direct selective expression of either E2Crimson or the reporter ChR2-tdTomato and Syp-EGFP. The insertion of a 13.7-kb spacer sequence in (B’) lowers the probability of F3 recombination.

### Optimal sparse labelling for AVP neuron visualization by SPARCL mice

AVP neurons represent a major neuronal subtype in the SCN, comprising approximately 4,000 cells, corresponding to about 20% of the total population of SCN neurons^19^. This suggests that a labeling ratio of about 0.025% (1 in 4000) may be ideal for visualization of these neurons; yet, consistently achieving such a low-density labeling has been an unfilled opportunity to date.

To test the sparseness performance of our genetic SPARCL system, we applied this system to imaging AVP neurons. We used an Avp-Cre transgenic mouse line (MGI:5697941) in which Cre recombinase is specifically expressed in AVP neurons^20,21^. To investigate our system, we first quantified the extent of sparseness of labelled AVP neurons in the SCN (Fig. 2) using two options of ChR2-tdTomato/Syp-EGFP lines (Fig. 1B and B’). As such, we prepared two mouse lines: AVP-Cre; SPARCL-FLPe^KI^; SPARCL-ChR2-tdTomato/Syp-EGFP^KI^ (2.6-kb inter-F3 distance) and AVP-Cre; SPARCL-FLPe^KI^; SPARCL-ChR2-tdTomato/Syp-EGFP^KI^ (16.3-kb inter-F3 distance) for sparseness test (see Fig. 2B and C).

**Figure 2.**
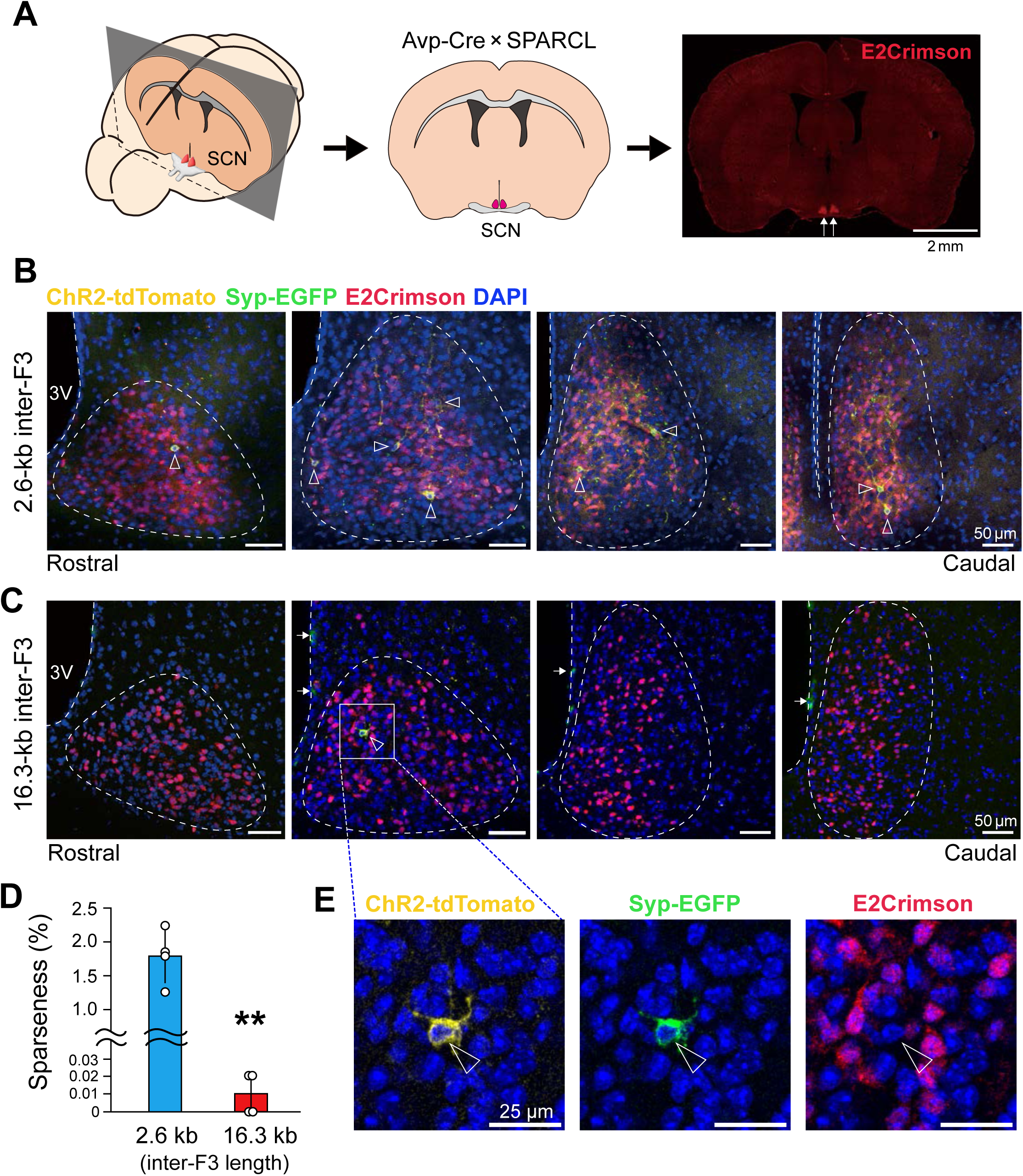
Sparseness performance of SPARCL for SCN AVP neurons. (A) Schematic brain histology illustrating the position of SCN. Avp-Cre × SPARCL mouse brain coronal section revealing E2Crimson-labelled AVP neurons in the SCN is provided. (B–C) Representative coronal brain sections of Avp-Cre × SPARCL mice bearing the inter-F3 sequence of 2.6 kb (B) or 16.3 kb (C). E2Crimson signals (red) represent negatively-selected majority of Avp-Cre⁺ neurons. In comparison, sparsely-labelled AVP neurons were illustrated by concomitant ChR2-tdTomato (yellow) and Syp-EGFP (green) signals (open arrowheads). The dashed lines delineate the shape of SCN in each section from the rostral to caudal sides. Non-specific anti-GFP signals were observed near the third ventricle (3V, white arrows). (D) Quantification of labeling sparseness in (B) and (C). Values are the percentage of ChR2-tdTomato⁺/Syp-EGFP⁺ double-positive cells in the total Avp-Cre⁺ neurons (mean ± s.d., *n* = 4 mice per group; \*\**P* < 0.005, two tailed t-test). (E) Magnified view of a singly labeled AVP neuron (arrowhead) from a 16.3-kb inter-F3 SPARCL mouse (corresponding to the boxed region in C). Images show ChR2-tdTomato (yellow), Syp-EGFP (green), and E2Crimson (red) signals merged with DAPI (blue). Scale bars, 25 µm.

A positively labelled AVP neuron(s) was surveyed across the SCN by immunohistochemical inspection of serial coronal brain sections encompassing the entire SCN; as shown in Fig. 2A, the SCN is located in the bottom of the hypothalamus and characterized by dense E2Crimson positive signals (*red*), consistent with AVP neurons being a major cell-subtype in the SCN. The distribution of AVP neurons in the SCN was also reflected by E2Crimson expression signals mainly located in the dorsomedial (shell) region of the SCN (Fig. 2B and C, *red*). In contrast and as expected, much fewer cells were found to exhibit positive reporter expression as revealed by sporadic distribution of ChR2-tdTomato (*yellow*); Syp-EGFP (*green*) co-expressing neurons in the SCN at the ratio of positively labelled cells to Cre^+^ total cells of 1.79 ± 0.40% for the 2.6-kb inter-F3 distance SPARCL and 0.01± 0.01% for the 16.3-kb inter-F3 distance SPARCL (Fig. 2 B,C, *n* = 4 mice for each group, mean ± s.d., *P* < 0.005, 2.6-kb versus 16.3-kb, Fig. 2D). These values correspond to an average of ∼70 (± 15) labelled cells per SCN for the former 2.6-kb and one or zero for the latter 16.3-kb SPARCL configuration. As shown in Fig. 2C and its enlarged view placed in Fig. 2E, a cell that exhibits robust coexpression of ChR2-tdTomato (*yellow*) and Syp-EGFP (*green*) was singly identified for the 16.3-kb configuration SPARCL mouse. This 1 or 0 minimal positive cell density per SCN appears excessively sparse, but it is still valid for individual cell tracing as it enables us to trace individual neurons without interference from neighboring cells. Therefore, we adopted the 16.3-kb line for all subsequent experiments.

### AVP neurons’ heterogenous morphology illustrated by SPARCL tracing

Our ultimate goal is to visualize the whole structure of AVP neurons; however, we recognized, during the course of our observations, that the SCN AVP neurons possess thin axons (<0.6 μm in diameter) that are hard to precisely trace using light-sheet fluorescence microscopy as well as the whole-brain serial sectioning tomography method, such as FAST^22^, at least in our current microscopy setting, providing an additional difficulty in tracing these AVP neurons. In our study, we thus applied two-photon scanning microscopy for imaging the SCN neurons at a higher resolution appropriate for tracing, albeit using a smaller brain-section (block) containing the whole SCN (see below) for the two-photon imaging — this condition cannot address our ultimate goal in full but we thought we might be able to provide a picture of previously uncaptured structures of AVP neurons and their heterogeneity in morphology in the SCN with the aid of our SPARCL mouse method.

We prepared a brain-section appropriate for imaging the structure of single AVP neuron inside and near-outside the SCN as a whole, using high resolution two-photon scanning microscopy. As illustrated in Fig. 3A, we prepared a pair of SCN hemisections, each approximately 3 mm AP × 2 mm DV × 0.5 mm ML in size, centered on a single-sided SCN, with the separation at the third-ventricle vertical plane (see **Methods** for cutting procedures). Although divided, this paired hemisections allowed tracing potential axonal connections between left and right SCN. The depth in ML direction (0.5 mm in thickness) was compatible with the working distance of high magnification, high numerical aperture two-photon micro-scanning. The brain blocks we prepared preserved the intact structure of the whole SCN and near-surrounding regions with no physical truncation inside the SCN. Using these block samples, we undertook two-photon scanning at a spatial resolution of 0.53 μm (*x*) × 0.53 μm (*y*) × 1.2 μm (*z*) and performed three-dimensional (3D) reconstruction of structure of each detected AVP neuron. The results obtained from 83 neurons were summarized in Table 1 and example axonal structures that were analyzed and categorized are presented in Fig. 3.

**Figure 3.**
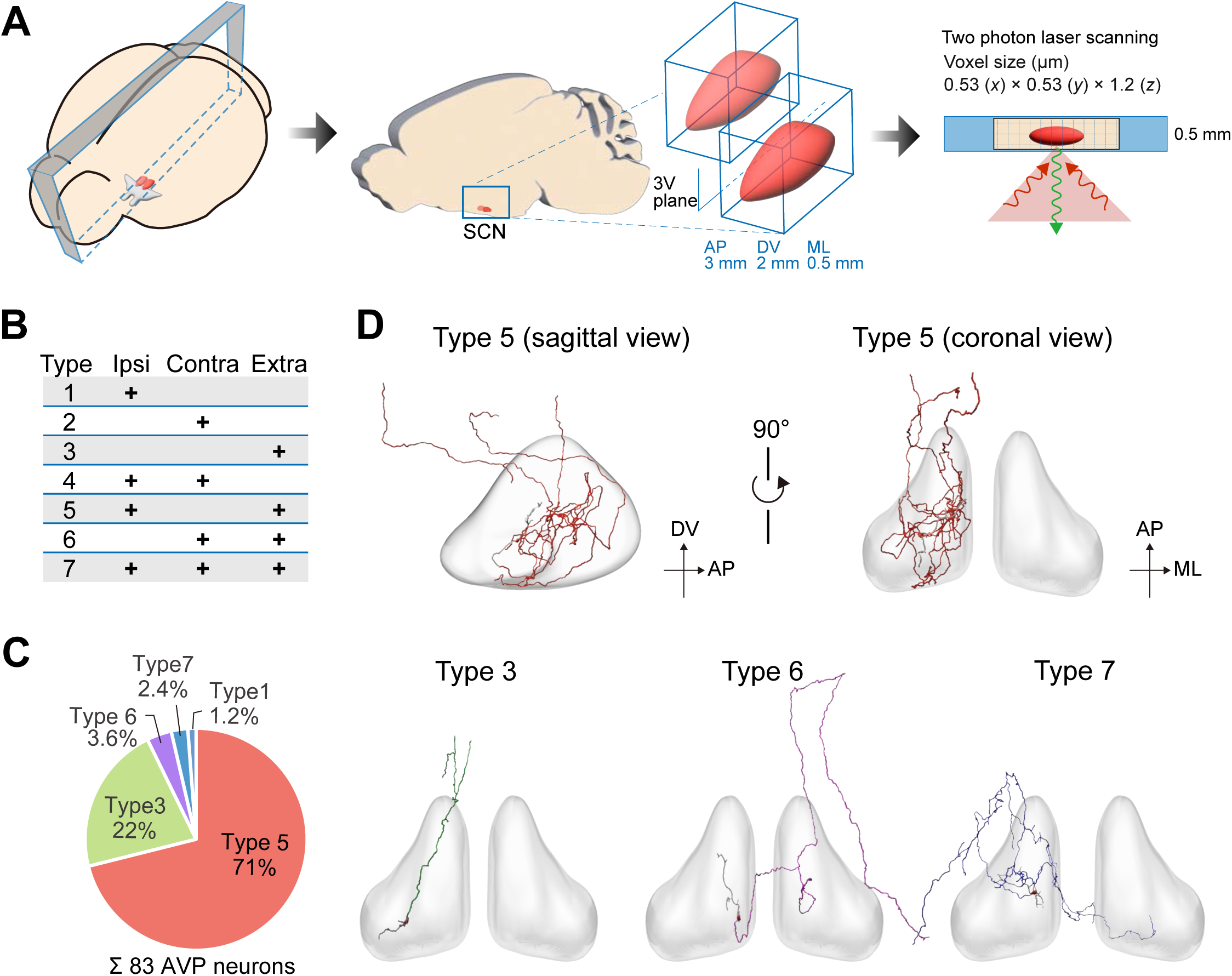
AVP neurons visualized by SPARCL in whole-mount SCN tissue imaging. (A) Experimental scheme of whole-mount SCN imaging. A sagittal brain section containing the whole SCN was divided into a pair of hemisections along the third ventricle (3V) and imaged using high-resolution two-photon microscopy. (B) Possible projection types based on the presence and absence of branches to ipsilateral (ipsi), contralateral (contra) and extra-SCN. (C) A pie chart showing the type prevalence of AVP neurons (SPARCL-identified, *n* = 83). (D) Representative axonal projection profiles of AVP neurons belonging to the types 3, 5, 6 and 7. The 3D SCN volume (gray) delineates the boundary of the SCN.

**Table 1.**
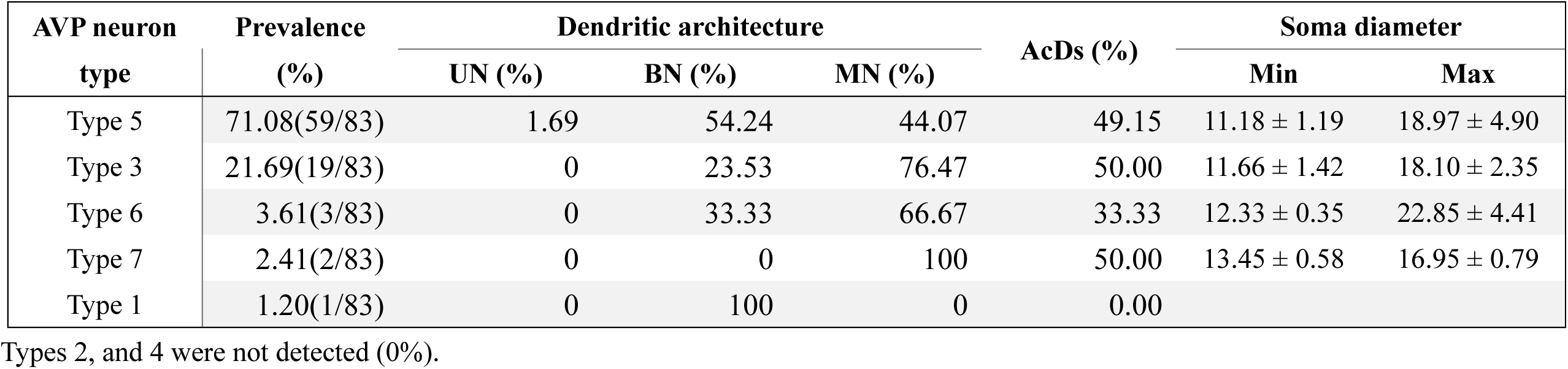

| AVP neuron<br>type | Prevalence<br>(%) | Dendritic architecture |  |  | AcDs (%) | Soma diameter |  |
| --- | --- | --- | --- | --- | --- | --- | --- |
|  |  | UN (%) | BN (%) | MN (%) |  | Min | Max |
| Type 5 | 71.08(59/83) | 1.69 | 54.24 | 44.07 | 49.15 | 11.18 ± 1.19 | 18.97 ± 4.90 |
| Type 3 | 21.69(19/83) | 0 | 23.53 | 76.47 | 50.00 | 11.66 ± 1.42 | 18.10 ± 2.35 |
| Type 6 | 3.61(3/83) | 0 | 33.33 | 66.67 | 33.33 | 12.33 ± 0.35 | 22.85 ± 4.41 |
| Type 7 | 2.41(2/83) | 0 | 0 | 100 | 50.00 | 13.45 ± 0.58 | 16.95 ± 0.79 |
| Type 1 | 1.20(1/83) | 0 | 100 | 0 | 0.00 |  |  |

Types 2, and 4 were not detected (0%).

A category list in Fig. 3B provides the possible maximum types of projection patterns of AVP neuron. For simplicity, we considered three potential axonal target zones: i) within the soma-resident (ipsilateral) SCN, ii) in the contralateral SCN, and iii) outside the SCN. Then, based on the presence or absence of axonal branch(es) (bifurcation/collateral) to i), ii) or/and iii), we considered seven possible combinatory patterns of axonal projections; which include those 1) only to the ipsilateral SCN, 2) only to the contralateral SCN, 3) only outside the SCN, 4) with dual target sites encompassing the ipsilateral and contralateral SCN, 5) both in the ipsilateral SCN and outside the SCN, 6) both in the contralateral SCN and outside the SCN, and 7) with targets encompassing all regions: the ipsilateral SCN, contralateral SCN and outside the SCN. On the basis of this framework, we classified the projection patterns of the 83 reconstructed AVP neurons (Table 1). In reality, five of the seven categories were observed as summarized in Fig. 3C. Among the five detected categories, the most dominant was type 5, representing approximately 71% of the total (59/83), followed by type 3, approximating 22%. Types 1, 6, and 7 were less common, only accounting for 1.2%, 3.6%, and 2.4%, respectively (Table 1). It is worth noting that four of the five detected categories shared a common feature: they all included projections outside the SCN (see representative structures of types 5, 3, 6, and 7 in Fig. 3D). In contrast, neurons displaying projections exclusively within the ipsilateral SCN were exceptionally rare, with only one such neuron identified among the 83 reconstructed neurons (Table 1: type 1, 1/83). Thus, “dedicated” local-circuit AVP neurons, if present as a subtype, do not appear to constitute a major structural class. These data indicate that, although local circuits are present in neurons of types 1, 5, and 7, the vast majority of AVP neurons with local ipsilateral projections also extend axons outside the SCN. This observation is consistent with the view that AVP neurons participate prominently in circadian output pathways. The examples of AVP neurons belonging to each type and their respective morphological characteristics are provided below in detail.

### Representative features of AVP neurons categorized based on projection patterns

We investigated potential heterogeneity or difference in structure between the types observed. Type 5, the most frequent form of AVP neurons, has axonal projections with branches ending in the ipsilateral SCN and extending to the extra-SCN (Fig. 4A). Type 5 neurons frequently show multiple intranuclear bifurcations and/or collaterals, with axons traversing large portions of the SCN (Fig. 4A, see axonal trajectories). In terms of dendritic structure, the majority (54%) exhibited two primary dendrites (Fig. 4C) classifying them as bi-dendritic neurons (BNs)^23^ (see also Table 1). About half (49.15%) of the neurons displayed axon-carrying dendrites (AcDs)^24^ — dendrites from which an axon originates rather than from the perikaryon (Fig. 4D) — besides the other half displaying axons originating from the perikaryon. The density map of the cell body of type 5 neurons in the SCN show a relatively uniform distribution across the E2Crimson-positive (thus, AVP-Cre positive) area in the SCN (Fig. 4E), suggesting no skewed distribution of type 5 across the SCN shell region.

**Figure 4.**
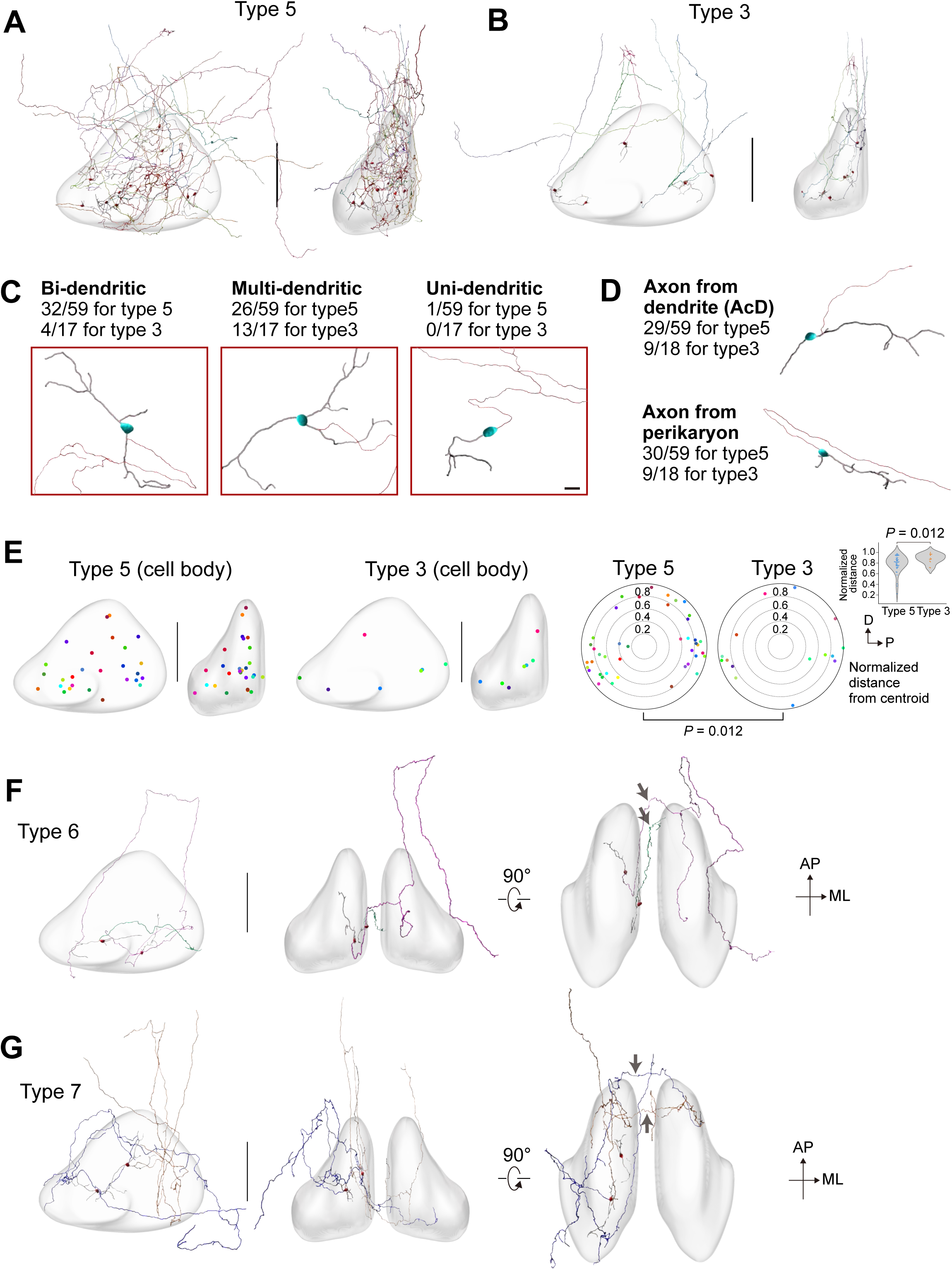
Topographic representation of AVP neuron structures in the SCN. (A) Topographic arrangements of type 5 AVP neurons in the SCN (*n* = 59, SPARCL identified). Colors represent different neurons. Although the SCN exists as a bilateral pair, reconstructed neurons from both sides were integrated into a common SCN reference space for presentation. (B) Topographic arrangements of type 3 AVP neurons in the SCN (*n* =18, SPARCL identified). (C) AVP neurons classified by the number of primary dendrites from the soma (uni-, bi- and multi-dendritic). Image examples show the structure of each category (soma, *blue*; axon, *red*; dendrites, *gray*) with the ratios in type 5 and type 3 AVP neurons. (D) AVP neurons classified by the site from which axon originates (AcD vs. perikaryon). (E) Soma distribution of type 5 and type 3 neurons within the SCN. *P* value was calculated using two-tailed Mann–Whitney U test. (F,G) Topographic arrangements of type 6 (F) and type 7 (G) neurons in the SCN. Arrows indicate the axons traversing from the ipsilateral SCN to the contralateral SCN through the posterior space between the SCN. AP, anterior-posterior axis. ML, medial-lateral axis.

Type 3 was the second most prevalent class in our dataset and projected exclusively outside the SCN without forming intra-SCN local circuit (Fig. 3C). Type 3 neurons can be regarded as dedicated “output” neurons; they have no intra-SCN branches, while forming arbors outside the E2Crimson-defined SCN (see axonal trajectories in Fig. 4B). To the best of our knowledge, AVP neurons with this projection type have not been reported previously; nevertheless, they represent approximately 22% of the total population of AVP neurons, indicating a previously unrecognized/overlooked subtype. With regard to dendrite architecture, the majority (76%) of type 3 exhibited more than three primary dendrites, classifying them as multi-dendritic neurons (MNs)^23^—a morphology suggesting a high capacity for synaptic integration; thus these neurons may act as an “integrator” to send output. Approximately 50% of the type 3 neurons displayed AcDs. The density map of the cell body of the type 3 neurons in the SCN led us to notice that they exhibit a preferential localization toward the periphery of the SCN, with a relative scarcity in the central region of the SCN (Fig. 4E, statistical evaluation of the location of neurons relative to the centroid of SCN, *p* = 0.012, type 3 vs. type 5).

Type 6 was rare in our dataset (*n* = 3), representing a type of neuron projecting toward the contralateral SCN (Fig. 4F). Type 6 is similar to type 3 in topology as both types provide projections exclusively outside the soma-resident (ipsilateral) SCN without forming intra-ipsilateral SCN local circuit. Type 6 is distinct, however, as its projections reach the contra-lateral (opposite) SCN (see axonal trajectories in Fig. 4F). Based on dendrite structure, the type 6 neuron that we observed was MN, bore AcDs, and the cell body was identified near the periphery of the SCN, similar to type 3.

Type 7 was also rare in our data (*n* = 2) and it displays projections to the contralateral SCN, like type 6, in addition to projections within the ipsilateral SCN and outside of the SCN. We emphasize that our finding of type 6 and type 7 neuron corroborates the presence of axonal projection bridging *left* and *right* SCN for AVP neuron (genetically defined AVP neuron), for which no specific studies addressed previously. Our data find that both type 6 and type 7 neuron get access to the contralateral SCN through a route beneath the third ventricle located between posterior regions of the SCN (see trajectories, Fig. 4F,G), which is compatible with a dense inter-SCN fiber connection reported in this area in previous studies^25^. As for dendrites, the observed type 7 neurons were classified as an MN, with its axon originating directly from the perikaryal. The cell body was positioned relatively centrally within the E2Crimson-positive SCN shell, similar to type 5.

Across the types, we did not observe marked differences in soma size (Table 1). Uni-dendritic neurons (UNs) were rarely observed, with only one such neuron identified in type 5. It should be, however, mentioned that the types 1, 6, and 7 were few in cell number, making it hard to eliminate problems of undersampling in concluding structural properties (UN, BN, vs. MN).

It is also our limitation that the numbers of bifurcations/collaterals shown in Table 1 refer to those formed inside the SCN: branches might exist outside the space of imaging, but we can not address this important issue in this study.

## DISCUSSION

While whole-brain projectome mapping remains an important long-term goal, we recognized, during the course of our investigations, that the SCN AVP neurons possess thin axons (see also refs^8,26,27^) that were hard to precisely resolve using light-sheet fluorescence microscopy as well as the whole-brain serial sectioning tomography method, such as FAST^22^, at least in our current microscopy setting, providing an additional hurdle in tracing SCN neurons besides obtaining the appropriate sparseness for labelling. Therefore, in our current study, we applied two-photon scanning microscopy for imaging the SCN neurons at a higher resolution, albeit using a smaller brain-section (block) containing the whole SCN — these conditions do not address our ultimate goal but have enabled us to reveal previously unseen structures of AVP neurons and their hidden heterogeneity in the SCN with the aid of our SPARCL method (cf. Figs. 3, 4 and Table 1).

Why has the sparse labelling of SCN neurons remained unexplored in previous studies? This is probably largely due to the unique anatomical situation of the SCN. Its location at the base of the hypothalamus, compact architecture (∼0.6 mm AP × ∼0.4 mm DV × ∼0.3 mm ML in size), densely packed organization of relatively small neurons totaling ∼10,000 cells in one side, and closely located left and right SCN with connections render this nucleus hard to sparsely label. We thought that controlled AAV infection (local/sparse/sporadic delivery of reporter by virus) may be impracticable for achieving consistent, low-density, specific labeling for SCN neuron. To address this inherent technical problem, we have devised hereby the GT-SPARCL — the genetic, two-step, sparse Cre/FLPe combination labelling method — that enabled labelling SCN AVP neurons approximately 2% when using AVP-Cre, SPARCL-FLPe^KI^ and SPARCL-ChR2-tdTomato/Syp-EGFP^KI^ (2.6-kb inter-F3 distance) in combination and 0.01% when using SPARCL-ChR2-tdTomato/Syp-EGFP^KI^ (16.3-kb inter-F3 distance; a longer variant) in lieu of the 2.6-kb version (see Figs. 1-2). To our knowledge, these are the first example of sparse labelling of genetically determined AVP neuron (i.e. AVP-Cre expressing neuron) in the SCN.

Based on the combination of axonal target zones, i.e., in the ipsilateral SCN, in the contra-lateral SCN, and/or outside the SCN, our SPARCL data identified five types of AVP-neurons, which we named as type 5 with projections to the ipsilateral SCN and outside the SCN, type 3 projecting exclusively outside the SCN, type 1 projecting only inside the ipsilateral SCN, type 6 projecting to the contralateral SCN in addition to those outside the SCN, and type 7 to all three regions. Furthermore, implying further potential diversity in structure, differential dendritic architectures (UN, BN, vs. MN) and/or axonal origins (AcDs or from perikaryon) were observed even in the same projection type (for example, in the type 5 neurons, 54% were BNs, 44% were MNs, 51% had axons from the perikaryon, and 49% had AcDs) (see Table 1). Thus, based on our SPARCL data, we conclude that SCN AVP neurons are not uniform— rather diverse— in view of morphology. Previous studies investigating the function of SCN AVP neurons using AVP-Cre mice did not take this diversity in AVP-neurons into account, forming a direction of future study. The diversity of cell characteristics among SCN AVP-neurons has also been recognized through single-cell RNA-seq data^28,29^; for example, Wen *et al.* found three AVP-neuron subtypes showing differential expression of *Vip, Nms, Cck,* and *C1ql3* (*Vip*⁺/*Nms*⁺/*Cck*⁻/*C1ql3*⁺, *Vip*⁻/*Nms*⁺/*Cck*⁻/*C1ql3*⁺, and *Vip*⁻/*Nms*⁺/*Cck*⁺/*C1ql3*⁺ neurons)^28^, while Xu *et al.* identified four different AVP-neuron subclusters using different set of genes involving *Syt10, Rorb, Vipr2,* and *Scg2* (*Syt10*⁺/*Rorb*⁻/*Vipr2*⁺/*Scg2*⁻, *Syt10*⁺/*Rorb*⁻/*Vipr2*⁺/ *Scg2*⁺, *Syt10*⁻/*Rorb*⁻/*Vipr2*⁺/*Scg2*⁻, and *Syt10*⁻/*Rorb*⁺/*Vipr2*⁻/*Scg2*⁻ neurons)^29^—These distinct transcriptional signatures may correspond to specific functions, and whether this molecular diversity aligns with the morphological heterogeneity identified in our study represents an important challenge requiring further investigation.

The prevalence of the most dominant type for AVP neurons, namely type 5 (neurons projecting to both the *ipsilateral* and *extra*-SCN), reached approximately 70%. This proportion is notably higher than previously reported. Pioneering work by Pennartz *et al.* in 1998 also described the presence of SCN AVP neurons possessing both local and extranuclear axons^14^; however, among the 83 reconstructed AVP neurons in their study, only a few were found to exhibit this pattern^14^. In their study, visually selected SCN neurons were filled with biocytin and immunostained to identify AVP^+^ cells within 200-µm-thick coronal sections^14^. The authors themselves noted that axons running beyond the plane of the slices were likely severed during sectioning, which could have led to an underestimation of extra-SCN projections. By contrast, our brain-block samples preserved the intact structure of the whole SCN and near-surrounding regions with no truncation inside the SCN. This allows more accurate estimation of the prevalence of the type 5 neurons in the SCN. Our data therefore revealed that the majority of AVP neurons (∼70%) residing in the SCN possess projections not only in the ipsilateral SCN but also extending to the extra-SCN. This notion is important in exploring the roles of AVP neurons in the SCN. It is tempting to speculate that most AVP neurons contribute to the intra-SCN network and circadian output signaling, simultaneously (and coordinately).

The second most predominant type in our dataset was type 3, which comprises about 20% of the AVP neurons examined. Type 3 formed projections exclusively outside the SCN without branches ending in the SCN. Interestingly enough, in previous studies, this type of neurons has been unrecognized (overlooked). This may not be a surprise, however. We found the location of type 3 neurons not ubiquitous across the SCN but rather skewed along the periphery of the SCN. Thus, these neurons may have escaped inclusion in previous studies, given that manual labelling approaches may result in unconscious selection of neurons that are located more centrally, suggesting the importance of non-biased approach. Additionally, the type 3 AVP-neurons identified by our study were mostly identified as MNs, i.e. neurons bearing more than 3 primary dendrites. This may help type 3 neurons act as integrators for circadian output signal.

In the present study, we also revealed for the first time the presence of commissural axons of AVP neurons (genetically determined AVP neurons) ending in the contralateral SCN. Van den Pol was the first to capture commissural fibers running between the left and right SCN in the horizontal SCN-tissue section labelled with HRP (horseradish peroxidase) via iontophoresis^8^, although their origin was unclear^8^. Via biocytin labelling in conjunction with AVP immuno-labelling, Pennartz *et al.* described that there exist AVP-ir fibers to the contralateral SCN^14^—however, the whole architectures of these AVP-ir neurons were not provided or pictured (as they used coronal sections). Intriguingly, Michel *et al.* showed that electrical stimulation of a single (unilateral) SCN evoked Ca²⁺ responses in the contralateral SCN in vitro, which were attenuated by pharmacological blockade of AVP receptors^30^— however, the structural entity of the connection was uncertain. In this context, our SPARCL data conferred a clear example of the whole-mount picture of the AVP neurons possessing commissural projections. Based on this whole-mount approach, we conclude that type 6 had simultaneous projections to both the contralateral SCN and extra-SCN, while type 7 had projections to all three regions, i.e. contralateral, ipsilateral, and extra-SCN. In our data, therefore, commissural projection-only neurons were not detected for AVP neurons (i.e. type 2) from our whole-mount approaches.

Furthermore, in our study, we detected a pathway (route) through which commissural fibers cross the midline. Interestingly, both type 6 and type 7 neurons accessed the contralateral SCN via a space between the posterior parts of the left and right SCN, running beneath the inferior margin of the third ventricle (see Fig. 3C) (also, note that in sagittal view, the SCN displays an elongated “fusiform” shape, lies in close vicinity to the optic chiasm for most of its length, but diverges gradually from the chiasm posteriorly to form this midline space^25^). We also observed that commissural fibers terminated in the shell of the contralateral SCN, which is compatible with earlier studies conducted by Leak and Moore, who suggested the presence of shell-to-shell bridging between left and right SCN by tracing retrograde multiple sequential trans-synaptic transport of pseudorabies virus (swine herpesvirus) injected to the dorsomedial hypothalamic nucleus (DMH, unilateral), a target site of the SCN (refs^31^). Our data thus extend the picture showing the presence of shell-to-shell connection to the single cell level resolution and in an AVP-neuron-defined manner. Our data suggest that neurons belonging to type 6 and type 7 may play a role in mediating bilateral SCN synchrony in addition to contributing to circadian output (both type 6 and type 7) and intra-SCN communication (only in type 7).

Finally, we considered it noteworthy that the vast majority of the AVP neurons reconstructed in this study possessed projections outside the SCN. Four of the five detected types—types 5, 3, 6, and 7—exhibited extra-SCN projections, whereas neurons projecting exclusively within the ipsilateral SCN were represented by only a single case in our dataset (type 1; 1/83). Thus, although a “dedicated” local-circuit AVP neuron was detected, it appears to be exceptionally rare and does not constitute a major subtype. Although micrographs provided by earlier studies often suggest the presence of likely local circuit neurons with all detected axonal ends confined in the SCN, it is possible that truncation of extranuclear projections may cause this implication. Indeed, the data from Moore *et al.* based on retrograde tracing of efferents from the SCN to DMH, SPZ, PVH, and PVT proposed the presence of extranuclear efferents from virtually all SCN neurons as they were labelled^31^. Our data are compatible with this previous model and suggest the important participation of each AVP neuron in extra-SCN outputs.

In conclusion, we reported here the structural heterogeneity and common feature of SCN AVP-neurons. In order to achieve stable sparse labelling of SCN neurons, we employed genetically-encoded two-step competitive (stochastic) recombination strategy, designated as GT-SPARCL, and applied it to visualizing AVP neurons in the SCN as a proof of concept of this sparse labelling method. Therefore, an intriguing next step will include extension of application beyond AVP neurons— GT-SPARCL may help to generate anatomical blueprints of different neuronal circuits defined by corresponding Cre lines, serving as a versatile tool for neuroscience. As a complementary method, our two-step strategy may contribute to the need for lower density (< 0.1%) cell labelling that has been difficult in one-step labelling approaches.

### Limitations of the study

An inherent limitation of this (our) and other sparse-labelling approaches is the sampling power imposed by low labelling density. Although sparse labelling at approximately 0.1% frequency enables high-resolution reconstruction of individual neurons and their long-range trajectories, it also requires large sample sizes to estimate population-level diversity with high statistical confidence. For example, if the target AVP neuron population consists of approximately 1,000 neurons, a 0.1% labelling probability would require on the order of 10,000 labelled observations to achieve near-complete sampling at a 95% confidence level. Obtaining such a dataset would require an impractically large number of genetically modified animals. Thus, although our analysis of 83 reconstructed neurons represents a substantial sampling effort for whole-mount sparse-labelling experiments, the possibility remains that rare morphological classes were not captured. This trade-off between morphological resolution and sampling completeness should be considered when interpreting our data.

## STAR METHODS

### RESOURCE AVAILABILITY

#### Lead contact

Further information and requests for resources and reagents should be directed to and will be fulfilled by the lead contact, Masao Doi.

### Material availability

The SPARCL mouse lines that we developed in this study have been deposited to the RIKEN BioResource Research Center (https://knowledge.brc.riken.jp/resource/animal/) under the foll-owing accession numbers: RBRC12840 for C57BL/6J Rosa26-SPARCL-13.7 kb spacer-ChR2-tdTomato-P2A-Syp-EGFP mice, RBRC12841 for C57BL/6J Rosa26-SPARCL-ChR2-tdTomato-P2A-Syp-EGFP mice, RBRC12842 for C57BL/6J Rosa26-SPARCL-FLPe mice.

All materials that were developed in this study are available from the lead contact with a completed Materials Transfer Agreement.

### Data and code availability

- All data reported in this paper will be shared by the lead contact upon request.
- This paper does not report original code.
- Any additional information required to reanalyze the data reported in this paper is available from the lead contact upon request.

## EXPERIMENTAL MODEL AND SUBJECT DETAILS

### Animals: SPARCL mouse lines

C57BL/6J Rosa26-SPARCL-FLPe knock-in (SPARCL-FLPe^KI^) and Rosa26-SPARCL-ChR2-tdTomato-P2A-Synaptophysin-EGFP knock-in (SPARCL-ChR2-tdTomato/Syp-EGFP^KI^) mice were generated at the University of Tsukuba. The SPARCL-FLPe^KI^ cassette was designed to include two Cre-dependent recombination unit pairs—loxP and lox3172—arranged in a cross-linked configuration. Positioned downstream of a 1.1-kb Rosa26 5′ homology arm and a CAG promoter, the first recombination-unit flanked by loxP sites consists of a lox3172 sequence and three copies of SV40 pA; this unit approximates 0.9 kb in size. The second recombination unit, which is flanked by lox3172 sites, encompasses the SV40pA(×3)-loxP sequence and the full-length E2Crimson coding-sequence followed by WPRE and SV40pA, totaling about 2.6 kb in size. Immediately downstream of the recombination units lies the FLPe coding sequence with a rabbit β-globin pA sequence followed by a 2.9-kb Rosa26 3′ homology arm. Similarly, the SPARCL-ChR2-tdTomato/Syp-EGFP^KI^ cassette includes two FLPe-dependent recombination pairs—FRT and F3—arranged in a cross-linked configuration. The first unit, flanked by FRT sites, contains an F3 and SV40pA(×3) (∼0.9 kb in total), while the second unit, flanked by F3 sites, includes the SV40pA(×3)-FRT and E2Crimson-WPRE-SV40pA cassette with or without a 13.7-kb spacer sequence for enhanced sparseness; this unit approximates either ∼2.6 kb or ∼16.3 kb depending on the spacer insertion. Immediately downstream of the recombination units lies the reporter cassette encoding ChR2(H134R)-tdTomato and Syp-EGFP linked via a self-cleaving P2A peptide. These SPARCL constructs were each inserted into the target Rosa26 locus in C57BL/6J zygotes by genome editing and site-specific integrase-mediated transgenesis as described^32,33^. Avp*-*Cre mice (Mouse Genome Informatics ID: 5697941)^20^ were crossed with a specific combination of SPARCL mice. Mice were reared on a regular 12-h light:12-h dark cycle with free access to food and water. All animal experiments were conducted in compliance with ethical regulations in Kyoto University and performed under protocols approved by the Institutional Animal Experiment Committee of the University of Tsukuba and the Animal Care and Experimentation Committee of Kyoto University.

## METHOD DETAILS

### Sparseness assessment by confocal immunofluorescence

For sparseness assessment, free-floating immunolabelling using 30-μm-thick serial coronal SCN sections was performed as described previously^34^ to intensify Syp-EGFP signal, using anti-GFP antibody (rat monoclonal, Nacalai Tesque, GF090R, 1:1000 dilution) and Alexa488-conjugated anti-IgG secondary antibody (Thermo Fisher Scientific, A11006, 1:1000 dilution). Sections were mounted in medium containing DAPI (Thermo Fisher Scientific) to stain nuclei. Fluorescence images were acquired using a confocal microscope (Leica SP8) with a ×40 oil-immersion objective lens. Excitation wavelengths were set as follows: 405 nm for DAPI, 488 nm for GFP and Alexa Fluor 488, 561 nm for tdTomato, and 594 nm for E2-Crimson. Emission spectra we recorded were 410-483 nm for DAPI, 500-550 nm for GFP and Alexa Fluor 488, 566-590 nm for tdTomato, and 615-784 nm for E2-Crimson. The number of cells expressing Syp-EGFP, ChR2-tdTomato, and E2-Crimson were counted using ImageJ software. Values were summed from the rostral to caudal margins of the SCN (∼20 sections per brain) for each animal. Sparseness was defined as the ratio of ChR2-tdTomato⁺-Syp-EGFP⁺ double-positive cells to all AVP fluorescent cells—that is the sum of ChR2-tdTomato⁺-Syp-EGFP⁺ double-positive cells and E2-Crimson⁺ cells, driven by Avp-Cre. The values (means ± s.d.) were determined using 4 independent animals.

### Whole-mount SCN tissue preparation

Animals aged from 6 to 12 weeks were selected for SCN fluorescence detection to minimize background autofluorescence signals. Before brain isolation, animals were anesthetized and subjected to transcardial perfusion using 4% paraformaldehyde (PFA) in phosphate-buffered saline (PBS). Following overnight postfixation in 4% PFA, isolated brains were embedded in 4% low-melting point agarose for sectioning using a vibrating microslicer (Neo-Linear Slicer NLS-AT, DSK). Sagittal 1,000-μm thick brain slices containing both sides of the SCN were cut out and these slices were trimmed under a microscope to excise a block centered on the SCN, approximately 3 mm along the anterior-posterior axis, 2 mm along the dorsoventral axis. Each block was then bisected along the third ventricle to yield a pair of 500 μm hemisections, each containing a single SCN. All trimming steps were performed in ice-cold PBS.

### Whole-mount SCN immunolabelling

The whole SCN hemisections (3× 2× 0.5 mm tissue block) were immunostained with anti-GFP antibody to obtain intensified Syp-EGFP signal. Before immunostaining, the sections were permeabilized with 0.5% Triton X-100 in PBS for 12 h and incubated in blocking buffer (PBS containing 3% bovine serum albumin and 0.5% Triton X-100) for additional 12 h. Then, the sections were incubated with anti-GFP antibody (Nacalai Tesque, GF090R, 1:1000 dilution) in blocking buffer for 24 h. Following extensive washes with 0.3% Triton X-100 in PBS (three times, 1 h each), the sections were stained with Alexa488-conjugated anti-IgG antibody in blocking buffer (Thermo Fisher Scientific, A11006, 1:1000 dilution) for 12 h. Thereafter, the sections were washed extensively (three times, 1 h each) with 0.3% Triton X-100 in PBS to get rid of free secondary antibody. All immunostaining procedures were performed at 4 °C.

Tissue clearing was subsequently performed using a modified SeeDB2G protocol^35^. We selected this tissue clearing method as it is known to cause minimal tissue swelling and neuron morphological deformation^36,37^, which is essential for our axonal tracing. Briefly, the tissue sections were incubated sequentially in 33%, 50%, and 97% Omnipaque 350 (GE Healthcare) in water with 0.5% Triton X-100 at room temperature, each for 1 h. Tissues were then immersed in 100% Omnipaque 350 for more than 2 h. The cleared tissues were mounted in 100% Omnipaque 350 between two coverslips separated by a 500-μm-thick silicone rubber spacer with a central cut-out, to preserve tissue architecture without causing compression.

### Neurite tracing by two-photon microscopy and 3D reconstruction

Tracing was performed using a two-photon excitation microscope (Olympus, FV1200MPE-IX83) with two GaAsP PMT detectors for EGFP (Alexa488) and tdTomato imaging. We used an Olympus ×30 silicone-immersion objective lens (UPLSAPO30XS, NA = 1.05, WD = 0.8 mm). EGFP (Alexa488) was excited at 930 nm and detected via a 495-540 nm bandpass filter. tdTomato (or E2Crimson) was excited at 1040 nm and detected using a 575-630 nm bandpass filter. Z-stack images were acquired at 1.2 μm intervals with an image resolution of 800 × 800 pixels and a zoom factor of 1.0. The brightness compensation function in the z direction was used to change detector sensitivity and laser power. A Kalman filter (line averaging = 2) was applied during data acquisition to improve signal-to-noise ratio.

Three-dimensional reconstruction of neurites was performed using Imaris software (ver. 10, Oxford Instruments). Using both ChR2-tdTomato and Syp-EGFP signals, neurites extending from a single soma were manually defined and 3D traced using the Filament Tracer module in Imaris. Neurites were classified as axons or dendrites based on projection length, shaft diameter, and bouton structure^8,19^. Axons were identified by their longer projections, relatively uniform and smaller diameters, and the presence of en passant boutons along the axon shafts as well as terminaux boutons at the terminal ends of axonal branches. In contrast, dendrites typically exhibited shorter and tapering processes with larger diameters and lacked axonal bouton structures. The 3D SCN volume was generated using the Surface module, in which we defined the boundaries of the SCN based on the distribution of E2Crimson cells, thereby providing a spatial reference for the reconstructed neurons and neurites. Soma surface was similarly generated using the Surface module using ChR2-tdTomato/Syp-EGFP signals, and soma diameter was estimated using the BoundingBoxOO Length parameter. The centroid of each SCN and soma was defined as the geometric center (center of mass assuming uniform voxel density) when analyzing the distribution of cell body of type 5 and type 3 neurons. To conduct comparison, each soma’s relative position was quantified as the distance from the soma centroid to the SCN centroid divided by the sum of its distance to the nearest SCN surface and that to the SCN centroid. Statistical difference was evaluated using a two-tailed Mann–Whitney U test due to the unequal sample size and skewed datapoint distribution.

## QUANTIFICATION AND STATISTICAL ANALYSIS

Cell counting for sparseness performance was quantified using ImageJ software. Statistical analyses and plots were generated with Python 3.9, using either a two-tailed Mann–Whitney U test or a two-tailed Student’s t-test.

## SUPPLEMENTAL INFORMATION

Supplemental information is available for this paper.

## ACKNOWLEDGEMENTS

We thank members of the M.D. laboratory, Drs Michiyuki Matsuda and Takahito Miyake for technical assistance and discussion. This work was supported in part by research grants from the Ministry of Education, Culture, Sports, Science and Technology of Japan (JP 22H04987, 23KF0246, 24H02306), the Basis for Supporting Innovative Drug Discovery and Life Science Research program of the Japan Agency for Medical Research and Development (JP21am0101 092), the Astellas Foundation for Research on Metabolic Disorders, the ABiS (JP16H06280, JP22H04926), the AdAMS (JP22H04922), and the Salt Science Research Foundation. This work was supported by Kyoto University Live Imaging Center. S.-W.H. is supported by a JSPS Postdoctoral Fellowship for Research in Japan.

## AUTHOR CONTRIBUTIONS

M.D., S.-W.H., Y.Y., and D.M. designed the research; S.-W.H., D.M., and S.D. performed experiments in collaboration with Y.Y., H.Z., S.M., S.T., H.Y., T.M., and E.H.; M.D. and S.-W.H. wrote the paper with input from all authors; M.D. supervised the project.

## DECLARATION OF INTERESTS

The authors declare no competing financial interests.

## Notes

### Competing Interest Statement

The authors have declared no competing interest.

